# A Pilot Study: Light-Based Estimation of Dim-Light Melatonin Onset Using a Wrist-Worn Illuminance Sensor

**DOI:** 10.64898/2026.11.26.769519

**Authors:** Akiyoshi Shimura, Kentaro Hayashi, Yuka Imaizumi, Junya Tanimoto

**Author notes:** Corresponding author: Akiyoshi Shimura, MD, PhD, Department of Somnology, Tokyo Medical University, 6-7-1 Nishi-Shinjuku, Shinjuku-ku, Tokyo 160-0023, Japan.

## Abstract

Dim-light melatonin onset is a reliable marker of human circadian phase, but its measurement requires repeated saliva or blood sampling under dim light. In this pilot study, we examined whether melatonin onset could be estimated from habitual light timing recorded by a wrist-worn sensor. Thirteen healthy adults wore illuminance sensors for 7 d before overnight salivary melatonin sampling in dim light (< 1 lx). Melatonin onset was determined with an individualized threshold, which identified onset reliably in all participants and was the main definition, and with a fixed 3 pg/mL threshold. Habitual light-onset time was computed at seven thresholds (1–300 lx), and the best-correlated threshold was selected post hoc. At 50 lx, light-onset time was closely associated with individualized-threshold melatonin onset (r = 0.924; 95% confidence interval, 0.76–0.98; slope, 1.035), which occurred 12 h 18 min ± 37 min later, but not with fixed-threshold onset (r = 0.146). Light-onset time remained significant after separate adjustment for self-reported wake time and actigraphy-derived mid-sleep. Root mean square errors were 41 min with leave-one-out and 63 min with nested cross-validation. These preliminary findings suggest that wrist-derived light timing may track melatonin onset and warrant larger studies with a prespecified threshold.

**Graphical Abstract:** 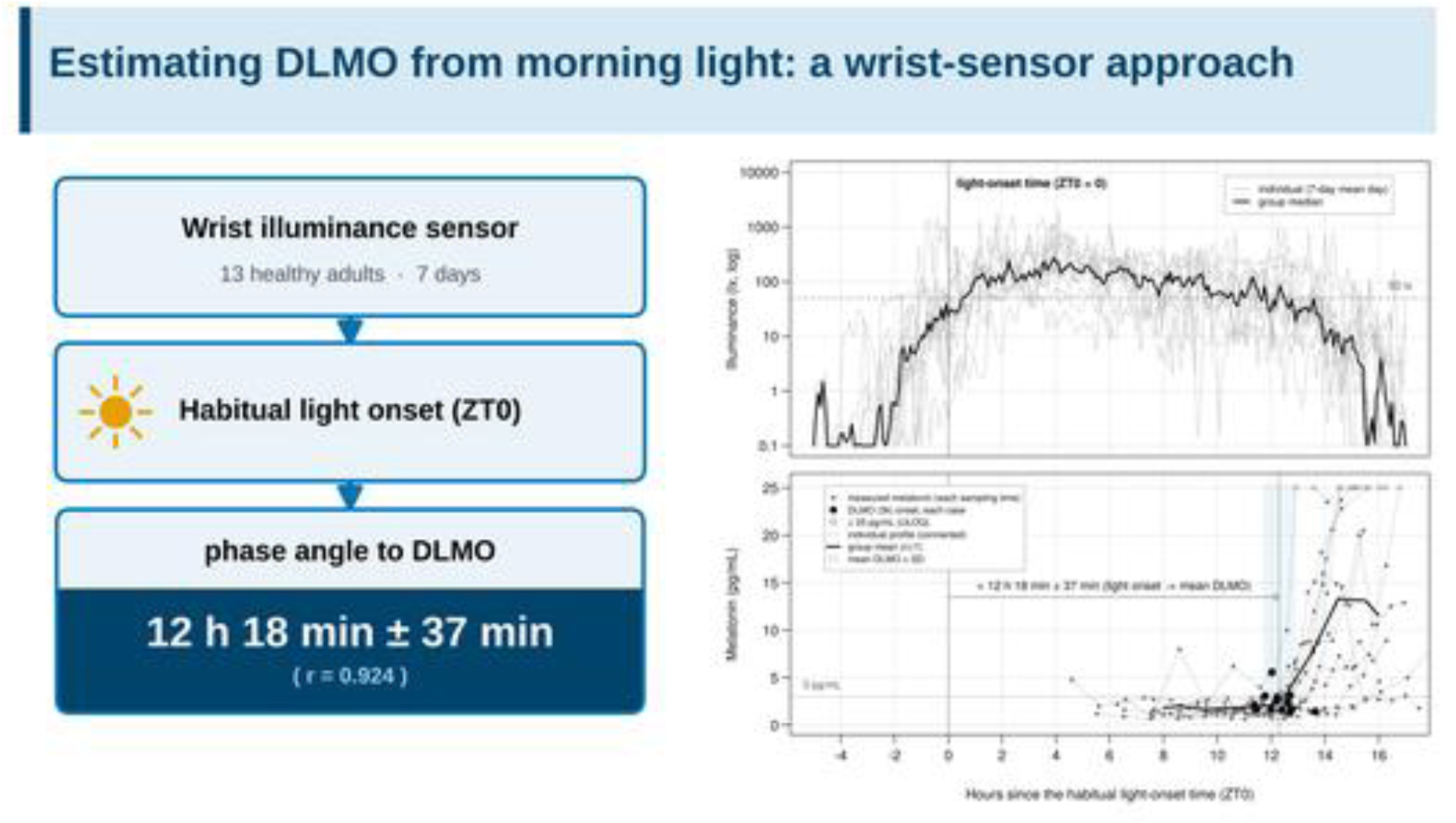

## Introduction

Melatonin, secreted by the pineal gland, is the principal hormonal signal of biological night and is often referred to as the “hormone of darkness.” Dim-light melatonin onset (DLMO), the time at which evening melatonin secretion begins under dim light, is one of the most reliable physiological markers of human circadian phase (Lewy and Sack 1989). Circadian phase varies markedly between individuals, and this variation is expressed in daily life as chronotype (Roenneberg et al. 2003, 2004). Because the phase response curve to exogenous melatonin is defined relative to DLMO, knowledge of an individual’s DLMO is essential for timing melatonin administration (Lewy et al. 1992). Circadian phase delay is also common in neurodevelopmental disorders such as autism spectrum disorder (Lorsung et al. 2021) and attention-deficit/hyperactivity disorder (Luu and Fabiano 2025), in which DLMO serves as the reference point for assessment and chronotherapy.

Despite this clinical importance, measuring DLMO imposes a substantial burden. Standard assessment requires repeated saliva or blood sampling, often at intervals of 60 min or less, over several hours under dim light; it is usually performed in a laboratory or clinical facility, although at-home protocols have been developed (Murray et al. 2024). The procedure is therefore costly and cumbersome, and results are not immediately available. In addition, the calculated DLMO varies with the sampling interval and the method used to set the threshold (Molina and Burgess 2011), and more robust approaches, such as the hockey-stick method, have been proposed as alternatives to fixed thresholds (Danilenko et al. 2014). This measurement burden remains a major barrier to the wider clinical use of DLMO and creates a need for a simple, physiologically interpretable estimate that could substitute for direct measurement.

Attempts to estimate DLMO more simply have therefore continued, but no definitive method has been established. Approaches reported to date (Stone et al. 2020) fall broadly into two groups. The first uses wearable sensor data, either by entering measured illuminance or activity into mathematical models of the circadian pacemaker (Stone et al. 2019; Cheng et al. 2021; Huang et al. 2021; Weed et al. 2026) or by feeding multivariate sensor features, such as light exposure, skin temperature, and activity, into machine-learning or regression models (Wan et al. 2021). The second estimates DLMO from sleep-timing measures such as bedtime, wake time, or mid-sleep (Martin and Eastman 2002; Burgess and Eastman 2005; Crowley et al. 2006). Most of these approaches require several variables or a fitted model, and where light is used, it is only one input among several. Model-based approaches may also require individual calibration, and their outputs can be difficult for clinicians to interpret or adjust. To our knowledge, no study has established a method that predicts DLMO from a single wearable measure of light exposure, without melatonin sampling or reliance on self-report.

Melatonin secretion is strongly regulated by light. Nocturnal light suppresses melatonin in a dose-dependent manner, particularly at short wavelengths (West et al. 2011); appropriately timed morning light advances the melatonin rhythm, including DLMO (Terman et al. 2001; Kozaki et al. 2011); and prior morning light can attenuate melatonin suppression by light later the same night (Kozaki et al. 2016). Beyond these regulatory effects, the rationale for estimating DLMO from light rests on the phase angle of entrainment: under stable entrainment, circadian phase maintains an approximately constant relationship with the timing of the light–dark cycle, and under natural conditions the clock entrains to the solar light–dark cycle such that DLMO occurs near sunset (Wright et al. 2013). The time at which habitual light exposure begins each day may therefore carry information closely related to individual DLMO and could serve as a practical proxy for melatonin-based circadian assessment.

We therefore hypothesized that the timing of an individual’s habitual light exposure is a key predictor of DLMO and that DLMO can be estimated from illuminance recorded continuously by a wrist-worn sensor in daily life. As a first step toward testing this hypothesis, we conducted a pilot study in which participants simultaneously wore two accelerometer–illuminance sensors, one on each wrist. Using salivary DLMO as the reference standard, we examined whether DLMO could be estimated from light timing alone, how this estimate compared with sleep timing and the core body temperature minimum, and whether light timing explained DLMO independently of self-reported wake time and chronotype. Because no illuminance threshold for defining habitual light onset has been established, a further aim of this preliminary study was to identify a candidate threshold and to estimate the likely predictive accuracy, as a basis for designing confirmatory studies.

## Materials and methods

### Participants and procedures

The study was conducted in Tokyo, Japan, and participants were recruited through the website of the collaborating institution and by direct invitation. Eligible participants were adults aged 20 y or older who could give written informed consent and who, if not engaged in shift work, had no marked variability in their sleep–wake schedule. Exclusion criteria were inability to stay overnight and undergo measurement under dim light, inability to wear the devices because of skin allergy or similar reasons, a sleep disorder other than a circadian rhythm sleep–wake disorder (circadian rhythm sleep–wake disorders, including shift work disorder, were not grounds for exclusion), and use of medications that could affect sleep or sleep-related measurements. Candidates were prescreened with a sleep diary and three questionnaires and were not enrolled if they screened positive on any questionnaire at its standard cutoff: the Pittsburgh Sleep Quality Index (global score > 5; Buysse et al. 1989; Doi et al. 2000), the Japanese version of the STOP-Bang questionnaire (score ≥ 3; Chung et al. 2012; Oshita et al. 2020), or the Center for Epidemiologic Studies Depression Scale (score ≥ 16; Radloff 1977).

Between January and February 2020, 23 volunteers gave consent and kept a sleep diary for 14 d, wearing two accelerometer–illuminance sensors, one on each wrist, during the second week. Ten were excluded because recording problems or poor wear compliance resulted in ≥ 2 d of missing data; the remaining 13, none of whom was engaged in shift work, stayed overnight on the final day for DLMO measurement. Because this was a pilot study, no a priori power calculation was performed; the target sample size was set with reference to previous studies that estimated circadian phase from multichannel ambulatory recordings under real-world conditions (n = 16 and n = 25; Kolodyazhniy et al. 2011, 2012). With 13 participants, the study had 80% power (two-sided α = 0.05) to detect correlations of approximately r ≥ 0.70.

The study was approved by the Ethics Committee of the Neuropsychiatric Research Institute (approval No. 184; October 30, 2019) and was conducted in accordance with the Declaration of Helsinki. All participants gave written informed consent before inclusion in the study and received a JPY 5,000 gift certificate for their time.

### Light exposure and sleep timing

Both sensors, the O:band (FAZOM / O: Inc.) and the Actiwatch Spectrum (Philips Respironics), recorded illuminance and activity at 1-min intervals. Sleep timing was obtained from the sleep diary, and sleep-corrected mid-sleep on free days (MSFsc) was calculated from the diary and interview data according to the Munich ChronoType Questionnaire (Roenneberg et al. 2003, 2004).

### Core body temperature

On the night of DLMO measurement, core body temperature was estimated continuously from forehead skin temperature with a zero-heat-flux monitoring system (3M Bair Hugger), which agrees closely with pulmonary artery temperature (Eshraghi et al. 2014); invasive measurements such as rectal temperature were not used. The core body temperature minimum (BTmin) was defined as the observed nadir when it was clearly identifiable and was otherwise estimated by cosinor fitting (Benloucif et al. 2005).

### Salivary melatonin and DLMO

Participants spent the measurement night in a light-controlled accommodation facility with blacked-out windows and remained in dim light (< 1 lx) throughout, consistent with current guidance on measuring melatonin in humans (Lockley 2020). Saliva was collected every 30–60 min, yielding approximately 10 samples per participant. Samples were stored frozen until shipment, and melatonin was measured by radioimmunoassay at a commercial clinical laboratory (SRL Inc., Tokyo, Japan). The upper limit of quantification was 25 pg/mL, and concentrations at or above this limit were reported as censored values (≥ 25 pg/mL).

DLMO was calculated using two definitions, neither of which was prespecified. In the individualized-threshold method (3k method, so named because the threshold is derived from the first three samples and a multiplier k), the threshold was the mean plus two standard deviations (SD; k = 2) of the first three samples obtained after the start of sampling, and DLMO was defined as the time at which melatonin first exceeded this threshold (Voultsios et al. 1997; Molina and Burgess 2011; Danilenko et al. 2014). In the fixed-threshold method (3pg method, named after its threshold), DLMO was defined as the time at which melatonin first exceeded 3 pg/mL (Murray et al. 2024); when baseline concentrations already exceeded 3 pg/mL, DLMO was taken as the time of the last upward crossing of 3 pg/mL after which concentrations remained above this level. In both methods, the time of threshold crossing was determined by linear interpolation between adjacent samples. Before any analysis involving light, the individual melatonin profiles were inspected, and the definition that identified DLMO reliably in all participants was designated as the main definition; the other definition was retained for comparison.

### Processing of illuminance data

Data from the two devices were aligned to a common 1-min time grid, and agreement between them was assessed as the Pearson correlation between their minute-by-minute illuminance series. From the illuminance (lx) recorded by each device, we computed the daily integrated illuminance; the temporal centroid of light exposure (the midpoint time of the daily integral of log illuminance); and, at thresholds of 1, 3, 10, 30, 50, 100, and 300 lx, the times of dark-period onset and dark-period offset (i.e., light-period onset), the dark-period midpoint, and the light-period midpoint. The dark period was defined as the longest continuous interval within a day during which illuminance remained below the given threshold; threshold crossings lasting 1 min or less were regarded as transient exposures or noise and ignored. Light-period onset was defined as the first time point after the dark period at which illuminance remained above the threshold for 2 min or longer, and the light period was defined as the remainder of the day. Daily values were averaged across all valid days of the 7-d recording period to yield each participant’s habitual value for each measure. The mean light-period onset time at the threshold selected in the analysis described below is referred to as the habitual light-onset time and is denoted ZT0, by analogy with zeitgeber time 0.

### Statistical analysis

Because this was a pilot study, three choices—the illuminance threshold, the DLMO definition, and the variables retained by backward elimination—were made within the same sample and are therefore exploratory; their implications are considered under Limitations. First, because no threshold for defining habitual light onset has been established, Pearson correlations between DLMO and light-period onset time were computed at each of the seven prespecified thresholds for both DLMO definitions. Light-period onset time at the threshold most strongly correlated with DLMO under the main definition was carried forward as the primary light measure. As a check for multiple testing across thresholds, a Bonferroni correction was applied to these 14 correlations. Second, as a supportive check of the DLMO definition, the variability (SD) of the phase angles between each DLMO estimate and two markers not derived from the illuminance recordings (BTmin and MSFsc; hereafter, light-independent markers, although both are themselves entrained by light) was compared with an F test, which had limited power at this sample size. Associations of each definition with the light measure are reported for completeness but were not used to select the definition.

Third, associations of DLMO under the main definition with five candidate predictors (age, MSFsc, self-reported weekday wake time, BTmin, and ZT0) and with the other light-timing measures at the selected threshold were examined with Pearson correlation coefficients and 95% confidence intervals (CIs) obtained by Fisher z transformation. The five predictors were entered into a multiple regression with backward elimination based on the Akaike information criterion. Because the number of predictors was large relative to the sample size, this model was complemented by partial correlations adjusting for one variable at a time and by hierarchical multiple regression (forced entry) with both orders of entry, which separated the incremental variance explained (ΔR²) by ZT0 and by wake time; the incremental contribution of adding MSFsc to the ZT0 model was evaluated in the same way. To assess robustness to measurement error in self-reported sleep timing, these analyses were repeated with an objective, actigraphy-derived mid-sleep time in place of self-reported wake time.

Predictive accuracy was assessed by leave-one-out (LOO) cross-validation of the simple regression of DLMO on ZT0 and summarized as the root mean square error (RMSE) and the proportions of estimates within ±1 and ±1.5 h. Agreement between measured and predicted DLMO was examined by Bland–Altman analysis, and Cook’s distance was used as an influence diagnostic. Because the threshold was selected using the full sample, this LOO estimate is potentially optimistic; we therefore also performed a nested cross-validation in which the threshold was reselected from the seven candidates within the training set of each fold, so that the held-out observation contributed to neither threshold selection nor model fitting. Associations with the amount and overall temporal distribution of light exposure (daily integrated illuminance and the temporal centroid of light exposure) were examined as secondary analyses. Data are presented as mean ± SD, and statistical significance was set at a two-sided p < 0.05. Analyses were performed and figures were created in R version 4.3.3 (R Foundation for Statistical Computing, Vienna, Austria; RRID:SCR_001905). This observational study is reported in accordance with the Strengthening the Reporting of Observational Studies in Epidemiology (STROBE) guideline.

## Results

The final analytic sample comprised 13 participants (five men and eight women; mean age, 31.9 ± 10.5 y). Mean self-reported wake times were 07:35h ± 1 h 30 min on weekdays and 09:35h ± 1 h 51 min on free days, and the weekly average sleep duration was 7.3 ± 0.4 h (Table 1). The minute-by-minute illuminance series recorded by the two simultaneously worn sensors were highly correlated (r = 0.993, p < 0.001).

**Table 1.** Baseline characteristics of participants.

| **Characteristic** | **Mean ± SD (n = 13)** |
|---|---|
| Sex, male / female (n) | 5 / 8 |
| Age (y) | 31.9 ± 10.5 |
| Bedtime, weekday<sup>a</sup> | 00:41 ± 1:35 |
| Bedtime, free day<sup>a</sup> | 01:26 ± 1:21 |
| Wake time, weekday<sup>a</sup> | 07:35 ± 1:30 |
| Wake time, free day<sup>a</sup> | 09:35 ± 1:51 |
| Sleep duration, weekly average (h) | 7.3 ± 0.4 |
| Mid-sleep on free days, uncorrected (MSF) | 05:30 ± 1:31 |
| MSFsc<sup>b</sup> | 05:03 ± 1:17 |
| Habitual light-onset time (ZT0)<sup>c</sup> | 09:36 ± 1:25 |
| Core body temperature minimum (BTmin) | 05:52 ± 1:02 |
| DLMO (3pg method) | 22:50 ± 1:16 |
| DLMO (3k method) | 21:54 ± 1:36 |

Values are mean ± SD (clock times in hh:mm; SDs in h:min) unless otherwise indicated. Twenty-three volunteers gave informed consent; 10 were excluded because of ≥ 2 d of missing data, yielding a final analytic sample of 13. <sup>a</sup>Self-reported. <sup>b</sup>Sleep-corrected mid-sleep on free days (Munich ChronoType Questionnaire). <sup>c</sup>7-d average, 50 lx threshold. BTmin, core body temperature minimum; DLMO, dim-light melatonin onset; MSF, mid-sleep on free days; MSFsc, sleep-corrected mid-sleep on free days; SD, standard deviation; ZT0, habitual light-onset time; 3k, individualized-threshold method; 3pg, fixed 3 pg/mL threshold method.

### Selection of the main DLMO definition

On inspection of the individual melatonin profiles, the fixed 3 pg/mL threshold could not reliably identify DLMO in two participants: in one, concentrations already exceeded 3 pg/mL at the start of sampling, so that DLMO could be determined only with the supplementary rule, and in the other, the threshold was exceeded at only a single sampling point (Supplementary Figure S1). The individualized threshold identified DLMO in all 13 participants. The 3k method was therefore used as the main definition in all subsequent analyses, and the 3pg method was retained for comparison.

### Threshold sensitivity and selection of the light measure

Pearson correlations between DLMO and light-period onset time at the seven thresholds are shown for both DLMO definitions in Table 2. For the 3k method, the correlation peaked at 50 lx (r = 0.924) and declined at both lower and higher thresholds, a nonmonotonic pattern; for the 3pg method, correlations were nonsignificant at all thresholds except 100 lx (r = 0.567; uncorrected p < 0.05). After Bonferroni correction for the 14 tests (α = 0.0036), only the 3k correlations at 30 lx (r = 0.780) and 50 lx (r = 0.924) remained significant. Light-period onset time at 50 lx was therefore carried forward as the primary light measure (ZT0).

**Table 2.** Pearson correlations between DLMO and light-period onset time defined at each illuminance threshold, for the 3k and 3pg methods.

| Illuminance threshold | 3k method | 3pg method |
|---|---|---|
| 1 lx | 0.505 | 0.250 |
| 3 lx | 0.613* | 0.265 |
| 10 lx | 0.730** | 0.206 |
| 30 lx | 0.780** | 0.168 |
| **50 lx<sup>a</sup>** | **0.924\*\*\*** | **0.146** |
| 100 lx | 0.559* | 0.567* |
| 300 lx (n = 9) | 0.298 | −0.429 |

n = 13 except as noted. Values are Pearson correlation coefficients between DLMO and light-period onset time defined at each illuminance threshold. \*p < 0.05, \*\*p < 0.01, \*\*\*p < 0.001 (uncorrected). <sup>a</sup>Threshold used to define the primary light measure. With Bonferroni correction for the 14 tests reported here (α = 0.05/14 = 0.0036), only the 3k correlations at 30 lx and 50 lx remained significant. DLMO, dim-light melatonin onset; 3k, individualized-threshold method; 3pg, fixed 3 pg/mL threshold method.

### Comparison of DLMO definitions (3k vs 3pg)

Mean DLMO was approximately 1 h earlier with the 3k method (21:54h ± 1 h 36 min) than with the 3pg method (22:50h ± 1 h 16 min), and the 3k method yielded a wider range of individual values (18:56h–00:26h vs 21:24h–01:54h; Table 3). For both light-independent markers, the phase angle was numerically less variable with the 3k method (SD of the phase angle to BTmin, 1 h 09 min vs 1 h 32 min; to MSFsc, 1 h 02 min vs 1 h 31 min), but the differences were not significant (BTmin: F(12, 12) = 1.76, p = 0.34; MSFsc: F(12, 12) = 2.18, p = 0.19).

**Table 3.** Comparison of two DLMO definitions (3k vs 3pg): phase angles with light-independent circadian markers and association with habitual light-onset time (ZT0)

| **Metric (mean ± SD unless otherwise indicated)** | **DLMO (3k)** | **DLMO (3pg)** |
|---|---|---|
| Mean DLMO (clock time) | 21:54 ± 1:36 | 22:50 ± 1:16 |
| DLMO range (earliest–latest) | 18:56–00:26 | 21:24–01:54 |
| Phase angle to BTmin | +7:59 ± 1:09 | +7:03 ± 1:32 |
| Phase angle to mid-sleep (MSFsc) | +7:10 ± 1:02 | +6:14 ± 1:31 |
| Phase angle from ZT0 to DLMO<sup>a</sup> | +12:18 ± 0:37 | +13:14 ± 1:46 |
| Correlation with ZT0, r (p) | 0.924 (<0.001) | 0.146 (0.64) |

n = 13. Mean DLMO and phase angles are mean ± SD (h:min). Phase angles to BTmin and MSFsc are the time intervals from DLMO to each marker (positive = marker follows DLMO); the phase angle from ZT0 is the interval from ZT0 to DLMO (positive = DLMO follows ZT0). <sup>a</sup>ZT0, habitual light-onset time (50 lx threshold; 7-d mean for each participant); the same measure is used in the correlation row. BTmin, core body temperature minimum; DLMO, dim-light melatonin onset; MSFsc, sleep-corrected mid-sleep on free days (Munich ChronoType Questionnaire); SD, standard deviation; 3k, individualized-threshold method; 3pg, fixed 3 pg/mL threshold method.

### Correlations between DLMO and candidate predictors

ZT0 showed the strongest correlation with DLMO of all candidate predictors (r = 0.924, p < 0.001; Figure 1 and Table 4). DLMO also correlated with the light-independent markers MSFsc (r = 0.765, p = 0.002) and BTmin (r = 0.690, p = 0.009), whereas its correlation with age was not significant (Table 4). Neither daily integrated illuminance (r = −0.178, p = 0.56) nor the temporal centroid of light exposure (r = 0.450, p = 0.12) was significantly correlated with DLMO.

**Figure 1.**
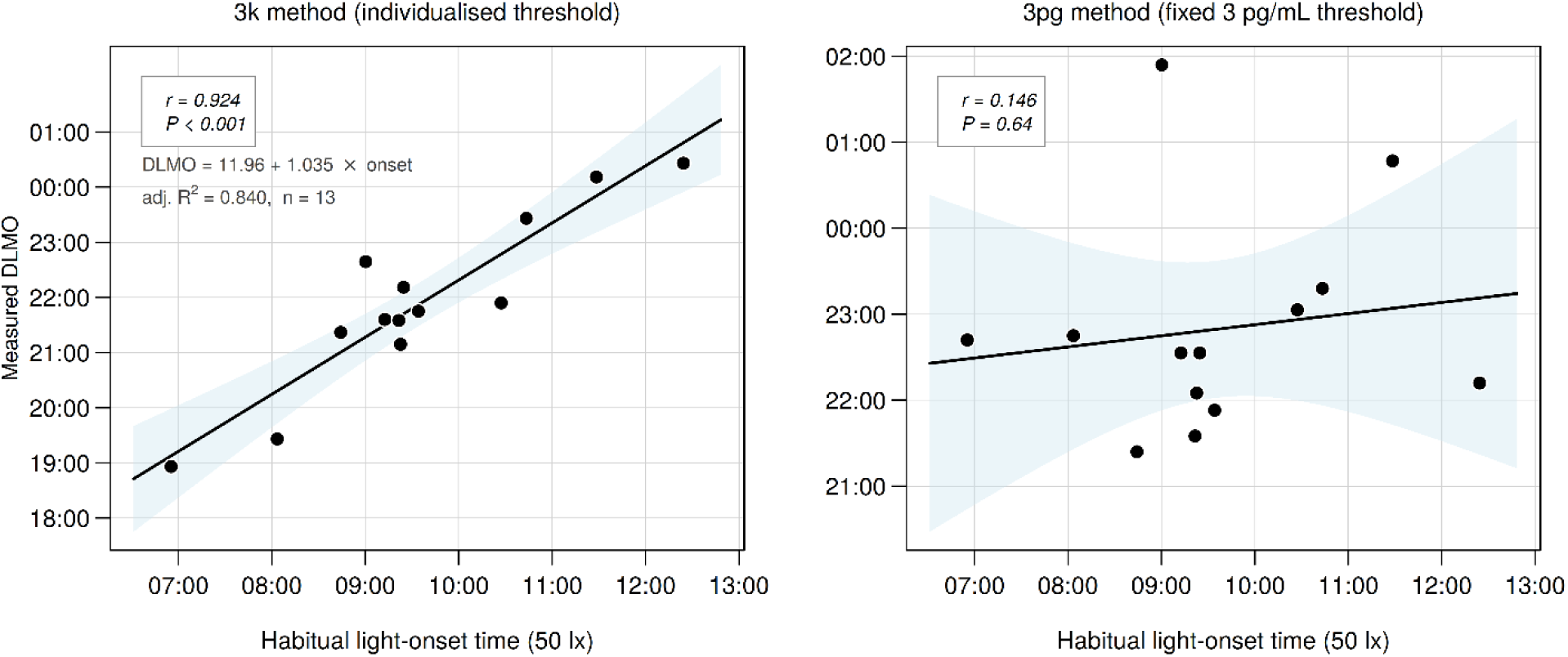
Association between habitual light-onset time (ZT0; 50 lx threshold) and measured DLMO under two DLMO definitions in 13 healthy adults. Left, individualized threshold (3k method; r = 0.924, p < 0.001); right, fixed 3 pg/mL threshold (3pg method; r = 0.146, p = 0.64). Each point represents one participant. Solid lines, least-squares regression lines; shaded areas, 95% confidence bands. The regression equation in the left panel expresses clock times in decimal hours. DLMO, dim-light melatonin onset; ZT0, habitual light-onset time; 3k, individualized-threshold method; 3pg, fixed 3 pg/mL threshold method.

**Table 4.** Associations of DLMO (3k) with candidate predictors and other light-timing measures: correlations and phase angles.

| **Predictor** | **r** | **p** | **Phase angle, mean ± SD (h:min)** |
|---|---|---|---|
| Age | −0.079 | 0.797 | — |
| MSFsc | 0.765 | 0.002 | +7:10 ± 1:02 |
| Core body temperature minimum (BTmin) | 0.690 | 0.009 | +7:59 ± 1:09 |
| Wake time, weekday<sup>a</sup> | 0.880 | <0.001 | +9:41 ± 0:46 |
| Dark-period onset (light offset)<sup>b</sup> | 0.416 | 0.158 | +1:19 ± 1:29 |
| Habitual light-onset time (ZT0)<sup>b</sup> | 0.924 | <0.001 | −12:18 ± 0:37 |
| Light-period midpoint<sup>b</sup> | 0.818 | <0.001 | −5:30 ± 0:57 |
| Dark-period midpoint<sup>b</sup> | 0.818 | <0.001 | +6:30 ± 0:57 |

n = 13. Pearson correlation coefficients. Light-timing measures are 7-d averages. Phase angle: time interval from DLMO (3k) to the predictor; positive = predictor follows DLMO, negative = predictor precedes DLMO. Thus the light-onset time precedes DLMO by 12 h 18 min and is shown here as −12:18; the same interval is expressed as +12:18 in Table 3, where it is measured in the opposite direction (from ZT0 to DLMO). <sup>a</sup>Self-reported. <sup>b</sup>50 lx threshold. BTmin, core body temperature minimum; DLMO, dim-light melatonin onset; MSFsc, sleep-corrected mid-sleep on free days; SD, standard deviation; ZT0, habitual light-onset time (50 lx); 3k, individualized-threshold method.

### Regression estimation of DLMO and phase angle

In simple regression, MSFsc and self-reported weekday wake time explained DLMO with adjusted R² values of 0.548 and 0.754, respectively. ZT0 showed the highest explanatory power (unstandardized regression coefficient B = 1.035; 95% CI, 0.750–1.321; adjusted R² = 0.840) and was the only variable retained when all five candidate predictors were entered into the multiple regression with backward elimination (Table 5).

**Table 5.** Multiple regression predicting DLMO (3k) with backward elimination.

| **Predictor** | **B** | **95% CI** | **β** | **t** | **p** |
|---|---|---|---|---|---|
| Intercept | 11.961 | 9.197–14.726 | — | 9.523 | <0.001 |
| Habitual light-onset time (ZT0) | 1.035 | 0.750–1.321 | 0.924 | 7.986 | <0.001 |

n = 13. Backward-elimination multiple regression predicting DLMO (3k). The candidate predictors entered were the same five variables examined by correlation (age, MSFsc, self-reported weekday wake time, the core body temperature minimum, and habitual light-onset time [ZT0, 50 lx]); backward elimination by the Akaike information criterion retained habitual light-onset time as the only predictor. Because the number of predictors was large relative to the sample size, this model is exploratory. Clock times are expressed in decimal hours, with times after midnight coded as values greater than 24. Final model: F(1, 11) = 63.78, p < 0.001, R² = 0.853, adjusted R² = 0.840. DLMO, dim-light melatonin onset; B, unstandardized regression coefficient; 95% CI, 95% confidence interval of B; β, standardized coefficient; t, t statistic; MSFsc, sleep-corrected mid-sleep on free days; ZT0, habitual light-onset time.

The phase angle of entrainment from ZT0 to DLMO averaged 12 h 18 min ± 37 min (Tables 3 and 4). DLMO preceded BTmin by 7 h 59 min ± 1 h 09 min and MSFsc by 7 h 10 min ± 1 h 02 min. When each participant’s 7-d mean illuminance profile and salivary melatonin profile were plotted relative to ZT0, the profiles converged, and melatonin onset clustered approximately 12 h after light onset (Figure 2; individual profiles in Supplementary Figure S1). Because aligning profiles to a common reference itself reduces visual dispersion, Figure 2 illustrates the phase relationship rather than providing independent evidence for it.

**Figure 2.**
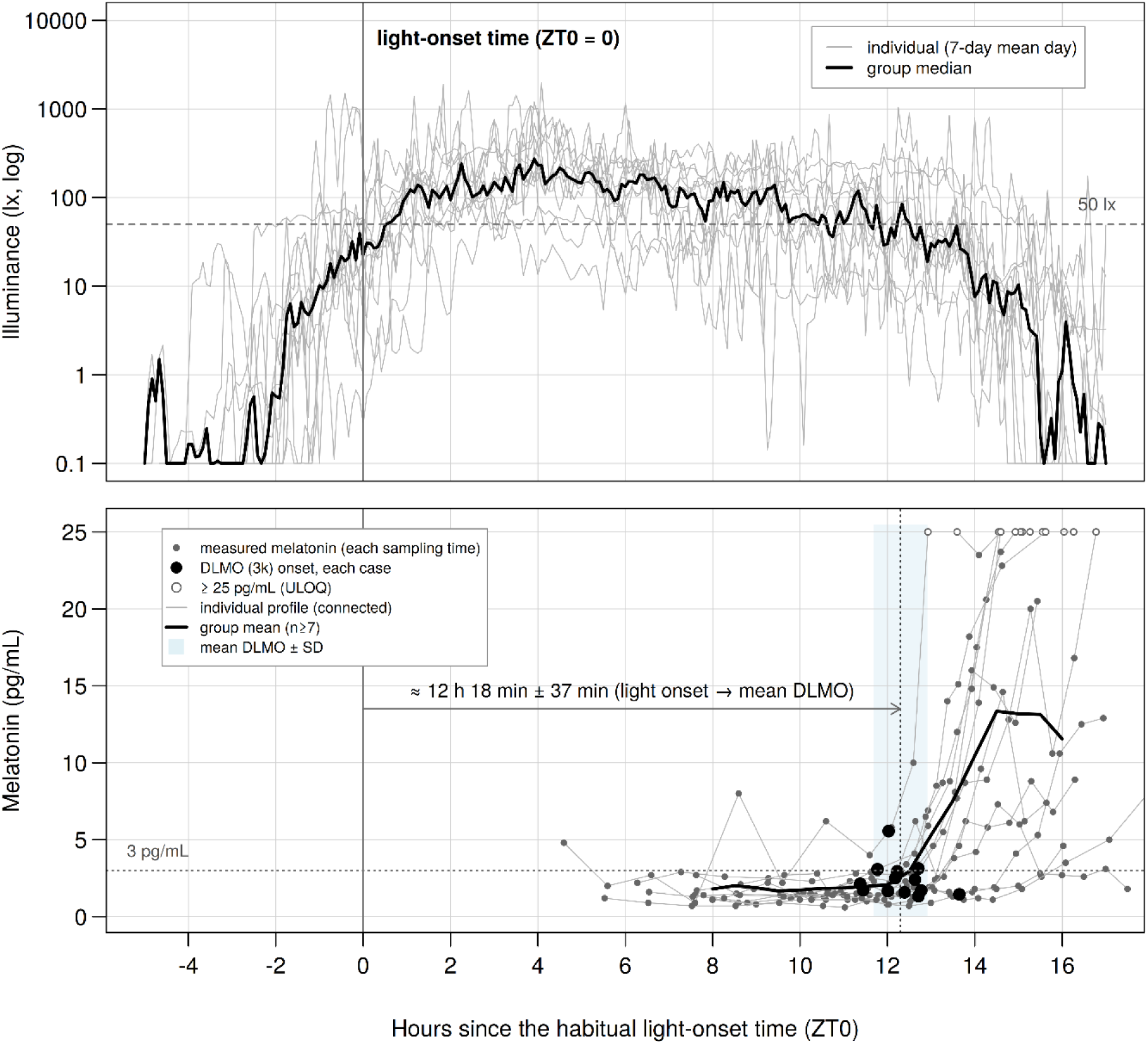
Light exposure and salivary melatonin in all 13 participants, aligned to each participant’s habitual light-onset time (ZT0 = 0). Upper panel: illuminance (logarithmic scale); thin gray lines, individual 7-d mean daily profiles; bold line, group median; dashed line, 50 lx. Lower panel: salivary melatonin; small gray filled circles, measured concentrations at each sampling time, connected by thin gray lines within each participant; bold line, group mean (shown only where at least seven participants contributed); large filled circles, individual DLMO (3k method); dotted vertical line and light-blue shaded band, mean ± SD of DLMO relative to ZT0; horizontal arrow, mean phase angle from ZT0 to DLMO (12 h 18 min ± 37 min); open circles, values at or above the upper limit of quantification of the assay (25 pg/mL); dotted horizontal line, 3 pg/mL. DLMO, dim-light melatonin onset; SD, standard deviation; ULOQ, upper limit of quantification; ZT0, habitual light-onset time (50 lx); 3k, individualized-threshold method.

### Adjustment for sleep timing

Self-reported weekday wake time correlated strongly with both DLMO and ZT0 (r = 0.880, p < 0.001 for both). When their contributions were separated, ZT0 made a significant independent contribution after adjustment for wake time (partial r = 0.661, p = 0.019; ΔR² = 0.098 for the addition of ZT0 to the wake-time model, p = 0.019), whereas wake time was not significant after adjustment for ZT0 (partial r = 0.372, p = 0.234; ΔR² = 0.020, p = 0.234). Because self-reported wake time is likely measured with greater error than the sensor-derived light measure, the analysis was repeated with actigraphy-derived mid-sleep time. The results were essentially unchanged: ZT0 remained a significant independent predictor after adjustment for objective mid-sleep (partial r = 0.636, p = 0.026; ΔR² = 0.098, p = 0.026), whereas objective mid-sleep was not (partial r = 0.146, p = 0.652; ΔR² = 0.003, p = 0.652). Adding MSFsc to the ZT0 model did not improve explanatory power (ΔR² = 0.004, p = 0.587).

### Predictive accuracy

The LOO-RMSE was 41 min, with 11 of 13 estimates (85%) within ±1 h and 12 of 13 (92%) within ±1.5 h. Bland–Altman analysis of the measured and LOO-predicted values showed a mean difference of −1.5 min, with 95% limits of agreement of −85 to +82 min (Figure 3). Cook’s distance was below the conventional cutoff (4/n = 0.31) for all participants (maximum, 0.25; Case 7 in Supplementary Figure S1), indicating that no single observation dominated the model. Because the 50 lx threshold was selected using the full sample, the LOO-RMSE is likely optimistic. In the nested cross-validation, in which the threshold was reselected within each training fold, the RMSE was 63 min (11 of 13 [85%] within ±1 h), and 50 lx was reselected in 12 of the 13 folds. In the remaining fold (Case 4), 30 lx was selected, and the prediction error for that case increased in magnitude from −63 min (LOO) to −182 min, which accounted for the entire difference between the two RMSEs.

**Figure 3.**
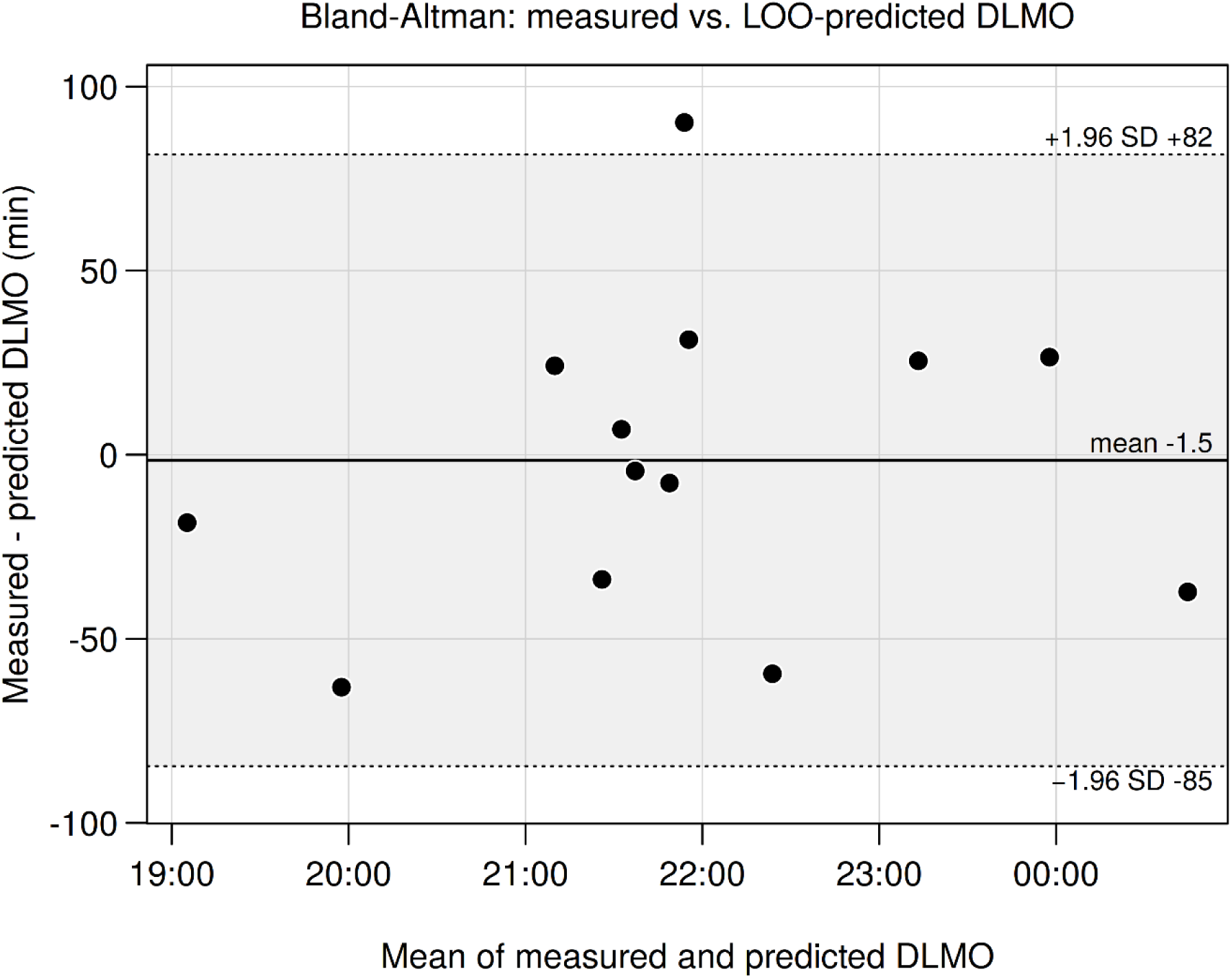
Bland–Altman plot comparing measured DLMO (3k method) with leave-one-out (LOO) cross-validated predictions from ZT0 (50 lx threshold) in 13 healthy adults. The horizontal axis shows the mean of the measured and predicted values as clock time; the vertical axis shows the difference (measured − predicted) in minutes. Each point represents one participant. Solid line, mean difference (−1.5 min); dotted lines, 95% limits of agreement (mean ± 1.96 SD; −85 to +82 min); shaded area, region between the limits of agreement. DLMO, dim-light melatonin onset; LOO, leave-one-out; SD, standard deviation; ZT0, habitual light-onset time.

## Discussion

In this pilot study of healthy adults, a single measure obtained with a wrist-worn illuminance sensor, ZT0, explained most of the interindividual variance in DLMO defined by the 3k method (r = 0.924; adjusted R² = 0.840). The regression coefficient (B = 1.035) was close to 1, indicating an approximately one-to-one correspondence in these data: a 1-h later ZT0 corresponded to an approximately 1-h later DLMO. If confirmed in independent samples, this relationship could serve as a simple rule for converting measured light timing into an approximate estimate of an individual’s DLMO. The association remained significant after adjustment for self-reported wake time (partial r = 0.661, p = 0.019), suggesting that the timing of light exposure may carry circadian phase information beyond that contained in the sleep schedule. Because it requires neither parameter estimation for a mathematical model nor multivariate learning, the approach may offer an interpretability and ease of implementation that model-based and multivariate approaches do not. These findings should nonetheless be regarded as preliminary, given the small sample and the exploratory selection of the threshold and DLMO definition.

As described in Results, the 3k method was adopted as the main DLMO estimate because the fixed 3 pg/mL threshold could not reliably identify DLMO in some participants: in one, baseline concentrations already exceeded the threshold, so that it did not separate the baseline from the circadian rise, and in another, it was exceeded at only a single sampling point. This choice was made on inspection of the melatonin profiles, before any analysis involving light. Because the 3k method uses each participant’s own baseline as the reference, it is not affected by the absolute level of secretion. Two further considerations support this choice. First, salivary melatonin concentrations are approximately 30% of plasma concentrations (Voultsios et al. 1997), so that in individuals with low secretion, measurement error near the lower limit of quantification has a relatively large influence; a method based on within-individual change may therefore be better suited to phase determination in saliva than one that depends on absolute concentration. Second, although the differences were not significant, the SDs of the phase angles to BTmin and MSFsc were smaller for 3k DLMO than for 3pg DLMO (Table 3), consistent with the known stability of the phase relationship between melatonin onset and core body temperature (Benloucif et al. 2005). Changing the definition from the 3k to the 3pg method markedly attenuated the association with ZT0 (r = 0.924 vs r = 0.146; Figure 1), and the corresponding 3pg-based regression model explained essentially none of the variance in DLMO (adjusted R² = 0.840 vs −0.068). This contrast was not used to select the definition, but it shows that the present findings depend strongly on how DLMO is defined.

Nevertheless, there is no international consensus on how DLMO should be calculated, and fixed-threshold methods have also shown clinical validity (Benloucif et al. 2008). Because they depend on absolute concentration, fixed-threshold methods are in principle susceptible to interindividual differences in baseline secretion, and the choice of method is known to affect estimated DLMO and comparability across studies (Glacet et al. 2023), although other reports indicate that its influence on circadian phase estimation is limited (Kennaway 2023). The instability of the 3pg method observed here is therefore unlikely to be a universal problem and more plausibly reflects the diverse secretion profiles of this small sample. Low secretors are relatively uncommon among healthy adults (Kennaway 2023), and their proportion in the present sample may reflect chance variation.

The nonmonotonic relationship between threshold and correlation has a plausible physiological and measurement basis. When the threshold was low (1–10 lx), light-period onset may have been identified prematurely and inconsistently because incidental dim light at night interrupted the dark period; when it was high (100–300 lx), onset was probably not detected until outdoor light levels were reached and was therefore recorded later than the biologically relevant transition. The 50 lx threshold lies within the range of illuminances at which melatonin suppression begins in many individuals (Phillips et al. 2019) and may reflect the first biologically significant light input to the circadian system. The correlation at this threshold may nonetheless have been limited by interindividual variability in light sensitivity (Phillips et al. 2019), which is further modulated by prior light history (Chang et al. 2011) and depends in part on age. The amount of light reaching the retina decreases with age owing to yellowing of the lens and senile miosis (Eto et al. 2020), and the magnitude of light-induced melatonin suppression differs markedly with age (Eto and Higuchi 2023): children show approximately twice the melatonin suppression of adults at comparable illuminances (Higuchi et al. 2014), whereas circadian photoreception is reduced in older adults (Turner and Mainster 2008). The optimal threshold and the absolute values of the phase angles obtained here are therefore likely to be sample-specific and may differ in environments dominated by natural light, where daytime light exposure is substantially higher (Wright et al. 2013). With n = 13, a contribution of noise to the nonmonotonic pattern itself cannot be excluded. Identifying a threshold that generalizes across individuals and settings is therefore a key objective for confirmatory studies.

In this sample, circadian phase was associated with the timing of light onset rather than with the total amount or overall temporal distribution of light exposure. Neither daily integrated illuminance (r = −0.178) nor the temporal centroid of light exposure (r = 0.450) correlated significantly with DLMO, whereas ZT0 was strongly associated with it. This is consistent with the principle that circadian entrainment depends critically on the circadian phase at which light acts on the pacemaker rather than on total light exposure alone (Duffy and Wright 2005) and suggests that, for phase estimation, the timing of light onset may be more informative than the total amount of light.

The approximately 12-h interval between ZT0 and DLMO also fits a broader physiological framework. The human phase response curve to melatonin is about 12 h out of phase with that to light (Lewy et al. 1998), such that light and melatonin act on the circadian system in an approximately antiphase manner. The close correspondence between habitual light onset and DLMO observed here is consistent with this framework and may offer an interpretable physiological basis for estimating melatonin phase from habitual light timing, although the exact interval depends on the illuminance threshold used to define light onset.

Wake time correlated strongly with DLMO (r = 0.880), but when the contributions of light timing and the sleep schedule were separated, the light measure retained significant independent predictive power, whereas the sleep-schedule variable did not. Because self-reported wake time is likely measured with greater error than the sensor-derived light-onset time, an asymmetry that could bias a partial correlation in favor of the light measure, we repeated the analysis with an objective, actigraphy-derived mid-sleep time. ZT0 remained a significant independent predictor (partial r = 0.636, p = 0.026), whereas objective mid-sleep was not (partial r = 0.146, p = 0.652), suggesting that the result is not merely an artifact of differential measurement error. Previous field studies are consistent with this interpretation: circadian phase advanced after a transition to natural light even though wake time remained essentially unchanged (Stothard et al. 2017), and DLMO advanced more than sleep onset (2.0 vs 1.2 h; Wright et al. 2013). Together, these observations support the possibility that light timing carries circadian phase information beyond that contained in the sleep schedule, although in the present sample the correlations of DLMO with ZT0 and with wake time were of similar magnitude (r = 0.924 vs 0.880), and this interpretation requires confirmation in larger cohorts.

These findings can be placed in the context of existing estimation methods. Wearable-based approaches fall broadly into two categories. The first enters wrist-measured illuminance or activity into mathematical models of the circadian pacemaker, such as limit-cycle or other differential-equation models. Such approaches have been validated in the field, including in shift workers (Stone et al. 2019; Cheng et al. 2021; Huang et al. 2021), and in ambulatory healthy adults, in whom a limit-cycle model driven by 9 d of light and activity data predicted DLMO with an error SD of approximately 1.1 h (Woelders et al. 2017); however, they require initialization and parameter estimation, and their accuracy depends on recording duration and light exposure (Weed et al. 2026). The second uses multivariate features, such as light exposure, skin temperature, and activity, in machine-learning or regression models to estimate DLMO directly (Wan et al. 2021). Such approaches require extensive melatonin data for training and may need recalibration to generalize across individuals, given interindividual differences in light sensitivity (Phillips et al. 2019). Estimation from the sleep schedule (Martin and Eastman 2002; Burgess and Eastman 2005; Crowley et al. 2006) relies on behavioral measures and is limited in accuracy: limits of agreement for estimates derived from questionnaires, sleep diaries, and actigraphy are approximately ±1.8 to ±1.9 h, and wake time alone correlates with DLMO at approximately r = 0.70 (Burgess and Eastman 2005; Reiter et al. 2020). In the present study, ZT0 alone gave an RMSE of 41 min with LOO and 63 min with nested cross-validation; we regard the latter, which also reselected the threshold, as the preferred, less optimistic estimate. However, wake time also correlated strongly with DLMO in this sample (r = 0.880), so part of the contrast with published sleep-based estimates may reflect characteristics of the sample, and direct comparison across studies is limited because populations, reference criteria, and validation schemes differ. What distinguishes the present approach is the suggestion that a single, directly measurable, and physiologically interpretable quantity, requiring neither a pacemaker model nor multivariate learning, may be sufficient, a possibility that warrants examination in larger studies.

This simplicity is particularly relevant to clinical implementation. Melatonin treatment timed by estimated DLMO has been reported to advance circadian phase similarly to treatment timed by measured DLMO (Swanson et al. 2024), suggesting that measured DLMO may not always be required in clinical practice and supporting the practical value of noninvasive estimates. If DLMO can be estimated from 7 d of illuminance recorded by a wearable sensor, outpatient assessment of circadian rhythm sleep–wake disorders and individualized timing of light and melatonin therapy could become more feasible.

### Limitations

This pilot study has several limitations. First, the sample was small (n = 13), and the 95% CI of the regression coefficient was relatively wide; the lower 95% confidence limit of the correlation (r = 0.76) approaches correlations reported for sleep-based estimates, so the apparent advantage of light timing should be interpreted cautiously. Participants were healthy urban adults studied in winter (January–February) under predominantly artificial lighting; generalizability to environments dominated by natural light, or to populations in which light exposure and sleep are dissociated, such as patients with circadian rhythm sleep–wake disorders or shift workers, is unknown. The study was conducted at a single site with a modest, self-selected sample that may not represent the full range of chronotypes. Second, the cross-sectional, observational design permits only associational inference; establishing whether light timing causally influences DLMO will require prospective and interventional studies. Third, confounding by the sleep schedule was assessed primarily with self-reported weekday wake time; although the principal finding was reproduced with actigraphy-derived mid-sleep, bedtime and sleep onset were not evaluated as separate confounders. Fourth, both the DLMO definition and the 50 lx threshold were selected within the same sample. The 3k method was adopted after inspection of the melatonin profiles rather than prespecified, and its advantage in phase-angle stability over the 3pg method was not significant (p = 0.19–0.34); the threshold was chosen post hoc because it yielded the highest correlation. Although the nested cross-validation, which reselected the threshold in each fold, gave a less optimistic RMSE of 63 min and reselected 50 lx in 12 of 13 folds, these choices remain sample-specific. Likewise, backward elimination with five candidate predictors in 13 participants is prone to overfitting and unstable variable selection, so the retention of ZT0 alone is exploratory, although partial-correlation and hierarchical analyses corroborated it. All principal findings are therefore restricted to 3k DLMO, and, as discussed above, a single 50 lx criterion may require adjustment across individuals, age groups, and lighting conditions. Fifth, DLMO was based on salivary melatonin, whose concentrations are lower and more variable than those in plasma (Kennaway 2024), and values at or above the upper limit of quantification (25 pg/mL) were censored. Sixth, illuminance was measured at the wrist and does not necessarily correspond to illuminance at the corneal plane, which is most relevant to retinal circadian photoreception (Lucas et al. 2014). Because the light reaching the eye varies with gaze direction and the spatial distribution of ambient light, spectacle- or head-mounted devices approximating corneal-plane illuminance would provide a more physiologically appropriate index. Light was also quantified as broadband illuminance rather than as melanopic equivalent daylight illuminance (mEDI), as defined in International Commission on Illumination standard CIE S 026 for responses mediated by intrinsically photosensitive retinal ganglion cells (Schlangen and Price 2021; Brown et al. 2022); the 50 lx threshold may therefore not transfer directly to light sources with different spectra. Because most commercially available wearables do not output mEDI, the fact that a broadband measure performed well here may facilitate practical implementation, although this requires confirmation against melanopic metrics. Seventh, device agreement was evaluated only as the correlation of the raw illuminance series, which are dominated by the day–night cycle; agreement at the level of derived light-onset times (for example, intraclass correlation or Bland–Altman analysis) was not assessed, and sensor calibration and wrist assignment were not evaluated. Eighth, light exposure was averaged over 7 d, whereas DLMO was measured on a single night, so day-to-day variability in light timing and any first-night effect could not be addressed; the menstrual-cycle phase of female participants was also neither controlled nor recorded, despite its potential influence on melatonin and temperature rhythms.

## Conclusions

In this pilot study of healthy adults, DLMO defined by the 3k method was closely associated with habitual light-onset time measured with a wrist-worn sensor (r = 0.924; 95% CI, 0.76–0.98; adjusted R² = 0.840), independently of self-reported wake time (partial r = 0.661) and objective mid-sleep (partial r = 0.636). DLMO occurred approximately 12 h after habitual light onset (12 h 18 min ± 37 min), and the regression coefficient was close to 1 (B = 1.035); this relationship could provide a simple heuristic for clinical use. Prediction errors of 41 min (LOO) and 63 min (nested cross-validation), together with LOO 95% limits of agreement of approximately ±1.4 h, indicate promising accuracy that is not yet sufficient for applications requiring precise phase timing. These preliminary results are specific to 3k DLMO in healthy adults under predominantly indoor, urban winter conditions, and the threshold was selected post hoc within the same sample. Preregistered external validation in larger, more diverse cohorts, with the analysis plan and illuminance threshold fixed in advance, is required before clinical application.

## Supporting information

Spl. fig 1

## Acknowledgments

The authors thank the study participants for their cooperation.

## Declaration of generative AI use

During the preparation of this manuscript, the authors used Claude (Anthropic), a generative AI tool, to assist with English-language editing, data processing, and writing the R code used to generate the figures. The authors reviewed and edited all output and take full responsibility for the content of this article.

## Data availability statement

The full dataset will be made publicly available upon acceptance of this article.

