## Supplementary material for "A Pilot Study: Light-Based Estimation of Dim-Light Melatonin Onset Using a Wrist-Worn Illuminance Sensor": Spl. fig 1

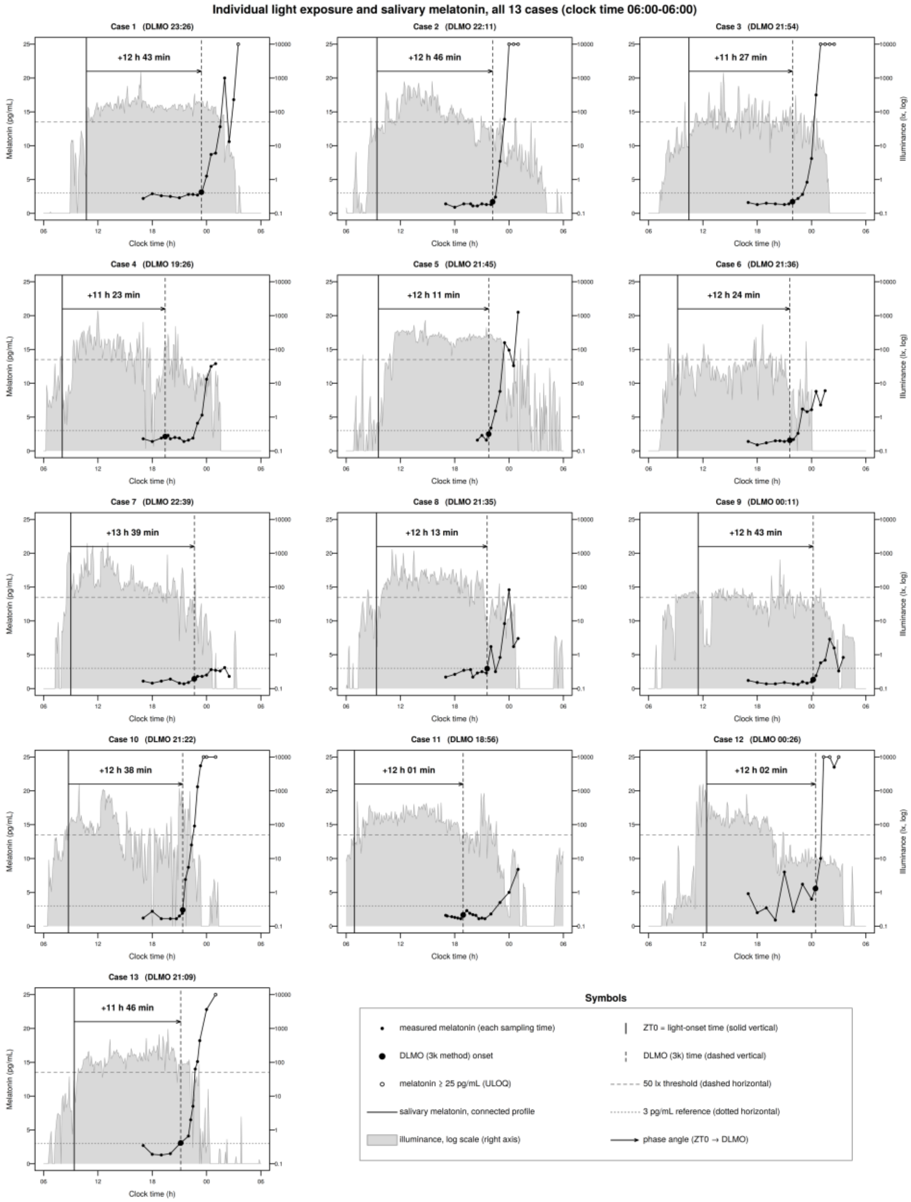


**Supplementary Figure S1.** Individual light exposure and salivary melatonin for all 13 participants (clock time, 06:00h–06:00h). In each panel: small black filled circles, the measured melatonin concentration at each saliva sampling time, connected by a black line (left axis); gray area, illuminance on a logarithmic scale (right axis); large filled circle, the DLMO (3k method); horizontal arrow (labeled with its value), the phase angle from ZT0 to DLMO; solid vertical line, ZT0; dashed vertical line, DLMO; dashed horizontal line, 50 lx; dotted horizontal line, 3 pg/mL; open circles, melatonin values at or above 25 pg/mL (upper limit of quantification). Panels correspond to Cases 1–13. DLMO, dim-light melatonin onset; ZT0, habitual light-onset time (50 lx); 3k, individualized-threshold method; ULOQ, upper limit of quantification.
